# A high throughput SEC23 functional interaction screen reveals a role for focal adhesion and extracellular matrix signalling in the regulation of COPII subunit SEC23A

**DOI:** 10.1101/2021.03.16.435679

**Authors:** Juan Jung, Muzamil Majid Khan, Jonathan Landry, Aliaksandr Halavatyi, Pedro Machado, Miriam Reiss, Rainer Pepperkok

## Abstract

Proteins that enter the secretory pathway are transported from their place of synthesis in the Endoplasmic Reticulum to the Golgi complex by COPII coated carriers. The components of the COPII transport machinery have been well characterized, but the network of proteins that regulate these components in response to extracellular cues have remained largely elusive. The discovery of such regulatory proteins is crucial for understanding how cells integrate extracellular and intracellular information to fine-tune membrane traffic in the secretory pathway. A key group of proteins that plays a central role in the regulation of membrane traffic is associated with the cytoskeleton. Using high throughput microscopy of the well-established VSVG transport from the ER to the plasma membrane, we comprehensively screened 378 cytoskeleton-associated proteins for their functional interaction with the COPII components SEC23A and SEC23B using a double knockdown approach. Ranking of the transport effectors identified 20 cytoskeleton-associated proteins as the strongest functional interactors of SEC23, most of them not previously associated with the secretory pathway. Knockdown of a subgroup of these interactors (FERMT2, MACF1, MAPK8IP2, NGEF, PIK3CA, ROCK1), associated with cell adhesion, not only induced changes in focal adhesions and the expression of adhesion-related genes but led to the specific downregulation of SEC23A, a SEC23 paralogue that has been implicated in the secretion of extracellular matrix (ECM) components. Furthermore, SEC23A downregulation could also be recapitulated by plating cells on ECM and was dependent on focal adhesions function. Altogether, our results identify a network of cytoskeleton-associated proteins connecting focal adhesions and ECM-related signalling with the gene expression of the COPII secretory machinery.

## INTRODUCTION

The secretory pathway is responsible for the delivery of proteins and lipids to their correct location, including the plasma membrane, extracellular space, and many membrane-bounded organelles. This allows the cells to dynamically regulate the size, composition, and function of those compartments. Alterations in the secretory pathway play important roles in many diseases including cancer, fibrosis, and neurodegenerative disorders (for reviews see (De Matteis and Luini 2011; Yarwood et al. 2020)), and understanding how cells integrate different internal and external stimuli to fine-tune secretion remains a central question in cell biology and disease mechanisms.

In the early secretory pathway, proteins are transported from their place of synthesis in the Endoplasmic Reticulum (ER) to the Golgi complex in a process mediated by COPII coated carriers. COPII assembly is initiated by the binding of the GTPase SAR1 to specific sites of the ER termed ER exit sites (ERES), followed by the recruitment of an inner coat composed of SEC23-SEC24 dimer responsible for cargo sorting and an outer coat composed of SEC13-SEC31 that provides a structural scaffold for membrane deformation and carrier formation (see (Miller and Schekman 2013) and references therein).

At ERES, protein secretion is regulated through an expanded COPII protein interaction network that ensures secretion meets physiological demands. For instance, nutrient deprivation reduces secretion by decreasing the number of ERES (Zacharogianni et al. 2011). Conversely, growth factor signalling increases the number of ERES and prepares the cells for a higher secretory demand (Farhan et al. 2010; Tillmann et al. 2015). More recently, Epidermal growth factor receptor (EGFR) signalling has also been linked to the transcriptional regulation of the COPII machinery in response to extended EGFR degradation to restore efficiently the availability of EGFR at the plasma membrane (Scharaw et al. 2016).

The cytoskeleton plays an important role in the organization and function of the secretory pathway by acting as a scaffold for membrane deformation, carrier transport, and organelle positioning (Anitei and Hoflack 2012; Gurel, Hatch, and Higgs 2014; Fourriere et al. 2020). For example, it has been shown that SEC23 interacts with DCTN1 (p150-glued), a subunit of the dynactin complex which links the COPII machinery to the microtubules, facilitating the long-distance movement of carriers towards the Golgi (Watson et al. 2005). A role of this SEC23-DCTN1 interaction in cargo concentration at ERES independent of microtubules has also been proposed (Verissimo et al. 2015). Less explored in the context of the regulation of the secretory pathway is the function of hundreds of cytoskeleton-associated and regulatory proteins that participate in the integration of extracellular and intracellular information (Moujaber and Stochaj 2019) including, for instance, mechanotransduction (Sun, Guo, and Fässler 2016), metabolism and signalling (Janmey 1998). To systematically and comprehensively explore the functional interactions of the cytoskeleton and associated factors with the transport machinery of the early secretory pathway, we conducted a functional siRNA-based interaction screen exploring possible synergistic interactions of 378 cytoskeleton and associated proteins with the COPII components SEC23A or SEC23B. Our results reveal an unexpected transcriptional regulation of the COPII subunit SEC23A by cytoskeleton-related proteins involved in focal adhesions and ECM function.

## RESULTS

### Identification of cytoskeleton-related proteins functionally interacting with SEC23A/B

Single siRNAs targeting altogether 378 cytoskeleton and associated proteins were co-transfected with siRNAs targeting SEC23A or SEC23B and biosynthetic transport of the vesicular stomatitis virus G protein (VSVG) from the ER to the cell surface (Kreis and Lodish 1986) was quantified as previously described (Simpson et al., 2012) (**Fig. 1A**). To identify synergistic functional interactors of SEC23A or SEC23B, the siRNA concentrations and transfection conditions were chosen such that transfection of SEC23A or SEC23B targeting siRNAs alone did not significantly affect VSVG transport compared to control transfected cells (**Fig. 1B**). However, co-transfection of siRNAs targeting SEC23A and SEC23B under these conditions caused significant transport inhibition of VSVG at the ER level (**Fig. 1B** and **Fig. 1E**) consistent with their role in cargo sorting and concentration at ERES.

**Figure 1.**
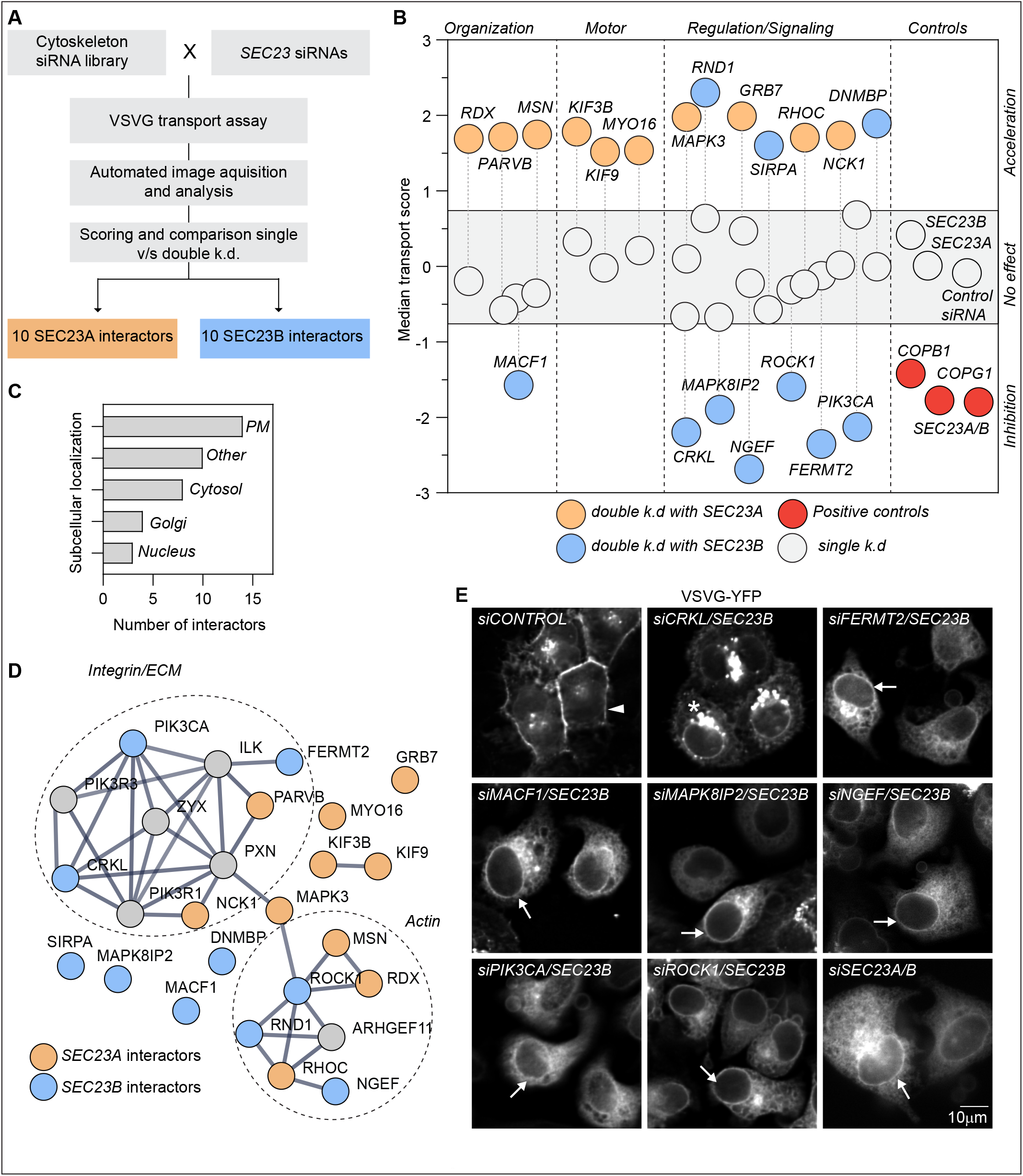
A functional interaction screen between SEC23 and the cytoskeleton-associated protein uncovers new proteins connected to secretion. **(A)** Screen workflow. HeLa cells were co-transfected with siRNAs targeting 378 cytoskeleton-associated proteins and siRNAs targeting the COPII subunits SEC23A or B. Using an automated microscopy acquisition and analysis pipeline, VSVG transport to the cell surface was quantified after 60 h of depletion in single and double knockdowns conditions. For each condition, a transport score was calculated. **(B)** Hit selection. The conditions in which the double knockdown score was significantly different from the expected additive effect of the single knockdowns were selected. Grey circles denote single knockdowns. Red circles denote the positive controls. Interactors in double knockdown are shown in orange (SEC23A interactors) or blue (SE23B interactors). **(C)** Annotation of the subcellular location of the interactors using the Human Protein Atlas and Compartments as the main sources. For proteins with multiple locations, only the two main locations were annotated (PM=Plasma membrane). *Others* include mitochondria, centrioles, focal adhesions, etc. **(D)** STRING network (Szklarczyk et al. 2019) using medium confidence and removing the text mining. Proteins in grey are not interactors but were included to fill up the gaps in the network. **(E)** VSVG transport assay. Widefield images of the VSVG-YFP after 1 h of temperature shift from 40°C to 32°C. Transport was assessed after 60 h of knockdown with the indicated siRNAs. Arrowhead indicates plasma membrane. Arrows indicate ER membranes. Asterisk indicates Golgi or Post-Golgi membranes.

Analysis of more than 85.000 images allowed us to rank the proteins targeted by the respective siRNAs according to their effect on VSVG transport when co-knocked down with either SEC23A or SEC23B (**Table 3**). Comparing transport rates of double knockdowns to single knockdowns and ranking of the strongest transport effectors revealed 20 cytoskeleton related genes in which VSVG transport in the double knockdown (SEC23A or SEC23B + cytoskeleton-associated protein) was significantly different from the additive effect one would expect from the results of the single knockdowns (**Fig. 1B**) (See material and methods for more details of the screen and data analysis).

The synergistic behavior in transport induced by the siRNA depletion strongly suggests a functional interaction between SEC23 and the 20 cytoskeleton related proteins selected by ranking, henceforth we call these proteins SEC23 functional interactors. We found SEC23 functional interactors that, in the double knockdown, led to transport acceleration, while others led to transport inhibition. Interestingly, VSVG transport inhibitors were only identified as functional interactors of SEC23B and no overlap between SEC23A and SEC23B interactors was observed (**Fig. 1B and Table 3**). The results of the screen are available in **Table 2**.

Annotation of the subcellular location using existing information in Compartments (Binder et al. 2014) and the Human Protein Atlas (Thul et al. 2017) shows that many of the interactors are proteins located at the plasma membrane with a few also localizing to the Cytosol, Nucleus or Golgi complex (**Fig. 1C**). Interestingly, none of the SEC23 functional interactors were annotated to localize to the ER or ER-exit sites as their main locations, which is the place of function for the COPII complex. While some of the interactors are motor proteins (KIF9, MYO16, KIF3B) or proteins that participate in the organization of the cytoskeleton (MACF1), most of the interactors are proteins involved in signalling pathways (**Fig. 1B**) that regulate the cytoskeleton in response to diverse stimuli including growth factor, actin signalling and cell-matrix interactions (**Fig. 1D**). Except for PIK3CA, ROCK1, and MACF1, these proteins have not been functionally implicated previously in the secretory pathway. Also, and in agreement with our previous genome-wide screen (Simpson et al. 2012), depletion of these proteins alone does not affect VSVG transport (**Fig. 1B**).

### Functional interactions at the ER exit level

For the double knockdown combinations causing inhibition of the VSVG transport, the intracellular location of VSVG was assessed, as the place of VSVG inhibition in the secretory pathway is expected to give first insights into the mechanism of how the respective target genes may functionally interact. A transport inhibition was observed when SEC23B siRNA was combined with siRNAs targeting any of these 7 proteins (CRKL, FERMT2, MACF1 MAPK8IP2, NGEF, PIK3CA, and ROCK1). For 6 of them (except for CRKL), VSVG was largely retained in the ER, similar to double knockdown of SEC23A and SEC23B, while in control transfected cells VSVG was observed mainly at the plasma membrane (**Fig. 1E**). Only for the double knockdown of SEC23B and CRKL, post-ER structures could be observed (**Fig. 1E**).

To gain more insight into the functional interactions we focused our analyses next on MACF1, as this protein has been previously also functionally implicated in the secretory pathway. MACF1 is a microtubule-actin cross-linker (Leung et al. 1999; Karakesisoglou, Yang, and Fuchs 2000) localized at focal adhesions and cell periphery but also present at the Golgi complex (Lin et al. 2005) where it has been proposed to be involved in Golgi complex to plasma membrane transport (Kakinuma et al. 2004; Burgo et al. 2012). To characterize the MACF1/SEC23B knockdown-induced transport impairment in more detail, the VSVG transport marker was released from the ER for 5 and 10 minutes only. In control cells, transfected with non-targeting siRNA, VSVG appeared in post-ER punctate structures visible already at 5 minutes after temperature shift, and a substantial amount of VSVG accumulated in the perinuclear region at 10 minutes, most likely representing the Golgi complex. In the case of MACF1/SEC23B double knockdown cells, occasional post-ER punctate structures became only visible at 10 min after the temperature shift and accumulation of the marker in the Golgi complex area apparently did not occur (**Fig. 2A**). Consistent with this transport impairment at the ER level upon MACF1/SEC23B double knockdown, we observed a reduction in the number of SEC31A positive ERES (**Fig. 2B, D)** and reduction of COPI positive ER to Golgi complex transport intermediates (**Fig. 2C, 2E**) compared to control transfected cells or cells transfected with the respective siRNAs alone. Moreover, P24, a cargo receptor that recycles between the Golgi complex and the ER, shifts its distribution to a more ER-predominant localization (**Fig. S1A)**. In MACF1/SEC23B double knockdown cells the Golgi marker GM130 appeared more dispersed compared to control transfected cells but not striking differences were observed (**Fig. S1B**). Also, no apparent changes were observed for the endosomal marker EEA1 (**Fig. S1C**). To confirm the transport inhibition observed for VSVG independently, we quantified the transport of E-Cadherin to the plasma membrane using the established inducible transport system RUSH (Boncompain et al. 2012). MACF1/SEC23B double knockdown showed a reduction of E-Cadherin transport between 40-70 %, depending on the siRNA used, while the single knockdowns did not show a significant transport alteration (**Fig. 2F, 2G**). In agreement with the data obtained for the transport marker VSVG, the majority of E-Cadherin was retained in the ER at time-points when in control transfected cells the transport marker already accumulated in post-ER structures resembling most likely the Golgi complex (**Fig. 2F**). Moreover, in the absence of any overexpressed cargo, electron microscopy analyses show that in MACF1/SEC23B double knockdown cells, the ER becomes often bloated, characteristic of a transport block at the ER level (see e.g. Fujiwara et al. 1988; Zhang et al. 1994) (**Fig. 2H**).

**Figure 2.**
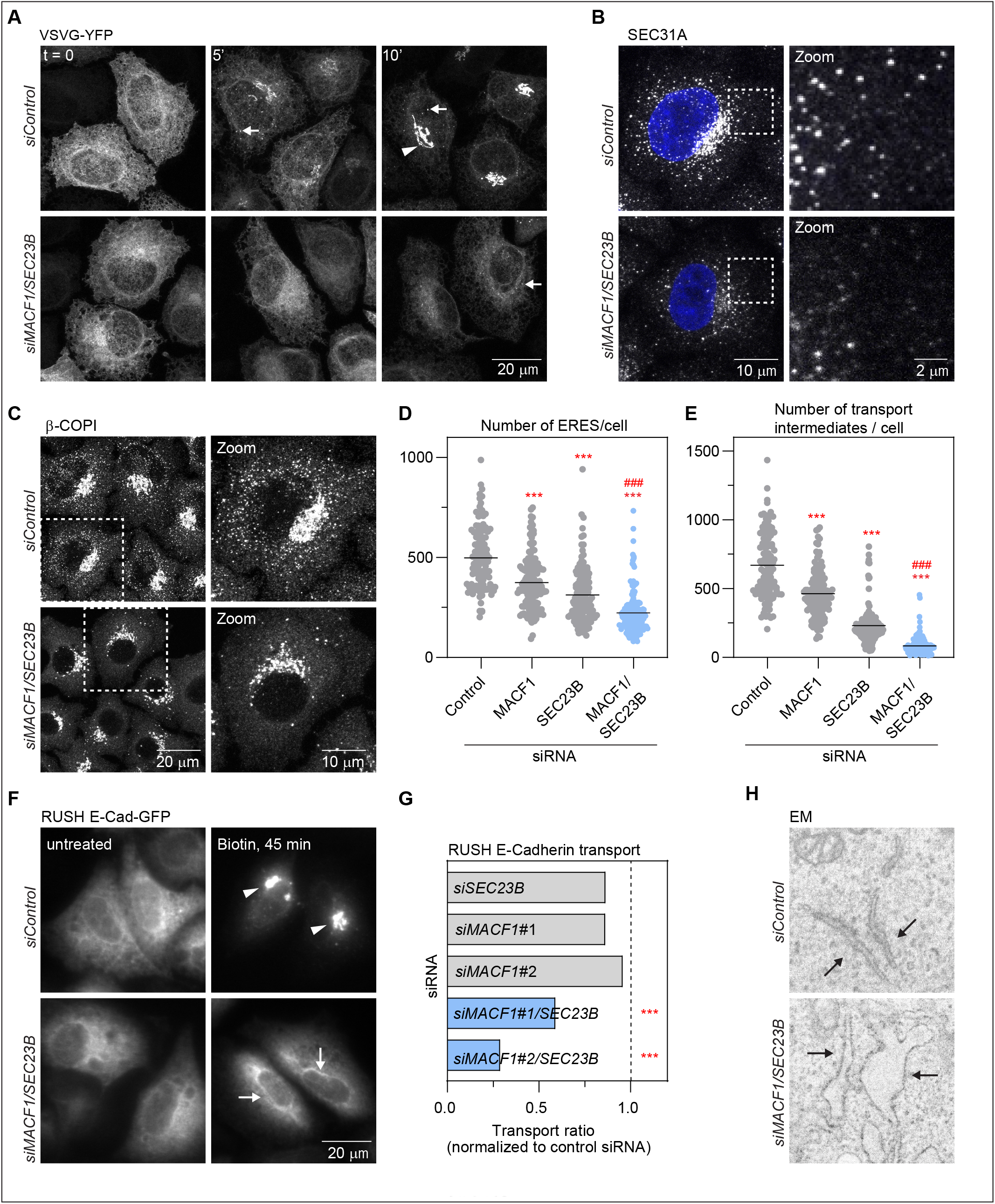
MACF1 and SEC23B co-depletion impairs ER to Golgi transport. **(A)** VSVG transport assay. Confocal images of VSVG-YFP at t=0 (40°C) and after 5 and 10 min of temperature shift from 40°C to 32°C. Transport was assessed after 60 h of knockdown with the indicated siRNAs. Arrows indicate puncta most likely representing transport intermediates. Arrowhead indicates juxtanuclear structures most likely representing Golgi. Confocal images of ERES assessed by SEC31A staining **(B)** and transport intermediated assessed by beta-COPI staining **(C)** after 48 h of knockdown with the indicated siRNAs. **(D** and **E)** Quantification of the images acquired in B and C respectively. **(F)** E-Cadherin RUSH transport assay. Widefield images of E-Cadherin-GFP at t=0 (untreated) and after 45 min of biotin release. The RUSH assay was performed after 48 h of knockdown with the indicated siRNAs. Arrows indicate ER membranes. Arrowheads indicate post-ER structures. **(G)** Quantification of the experiment shown in F. The bars represent the transport average of one representative experiment in which > 500 cells per condition were quantified and normalized to control siRNA. **(H)** Electron microscopy images of cells treated with the indicated siRNAs for 48 h. Arrows indicate ER membranes. The bloated ER in the MACF1/SEC23B double knockdown is characteristic of an ER transport block. Statistical significance: *** p<0.001 compared to control. ### p<0.001 compared to SEC23B or MACF1 single knockdowns, Student’s t-test.

Next, we asked whether MACF1 and SEC23B functional interaction is due to physical interaction. While most of the MACF1 staining is located in the cytosol, cell periphery, and Golgi (as previously described) with no significant staining in the ER or peripheral exit sites, we cannot discard colocalization with exit sites in the densely populated perinuclear region (**Fig. S1D**). To test if MACF1 and SEC23B interact physically we performed Co-IP experiments but we failed to detect an interaction under our experimental conditions (**Fig. S1E**). While these results do not exclude completely the possibility of physical interaction of SEC23B and MACF1 they suggest that the functional interaction of these two proteins as observed here is most likely indirect.

### Knockdown of specific functional interactors leads to SEC23A downregulation

The striking similarity of the VSVG transport block phenotype for 6 out of 7 SEC23B functional interactors in the double KD experiments suggests that there might be a common mechanism or pathway by which these proteins functionally interact with SEC23B. Based on their localization and functional annotation reported in the literature, we considered them unlikely to be direct physical interactors of SEC23B, although this possibility cannot be rigorously excluded. We speculated that their knockdown in cells may affect factors that are more directly related to the membrane traffic machinery in the early secretory pathway. As a first step to investigate this hypothesis, we performed mRNA sequencing to explore the transcriptional changes induced by depletion of these 7 interactors alone and testing if their depletion might cause overlapping transcriptional changes. With a p-adjusted value of 0.1 and a Log2 fold change above 0.58 or below −0.58, we found significant changes in gene expression for all the knockdowns tested (**Fig. 3A** and **Table 4** for the gene list). As expected, depletion of a protein with pleiotropic functions like PIK3CA generated the biggest changes (i.e. 419 genes downregulated and 580 genes upregulated). Also, out of the approx. 2300 genes differentially expressed across all 7 conditions tested, we observed little overlap between the transcriptional profiles (**Fig. 3B**).

**Figure 3.**
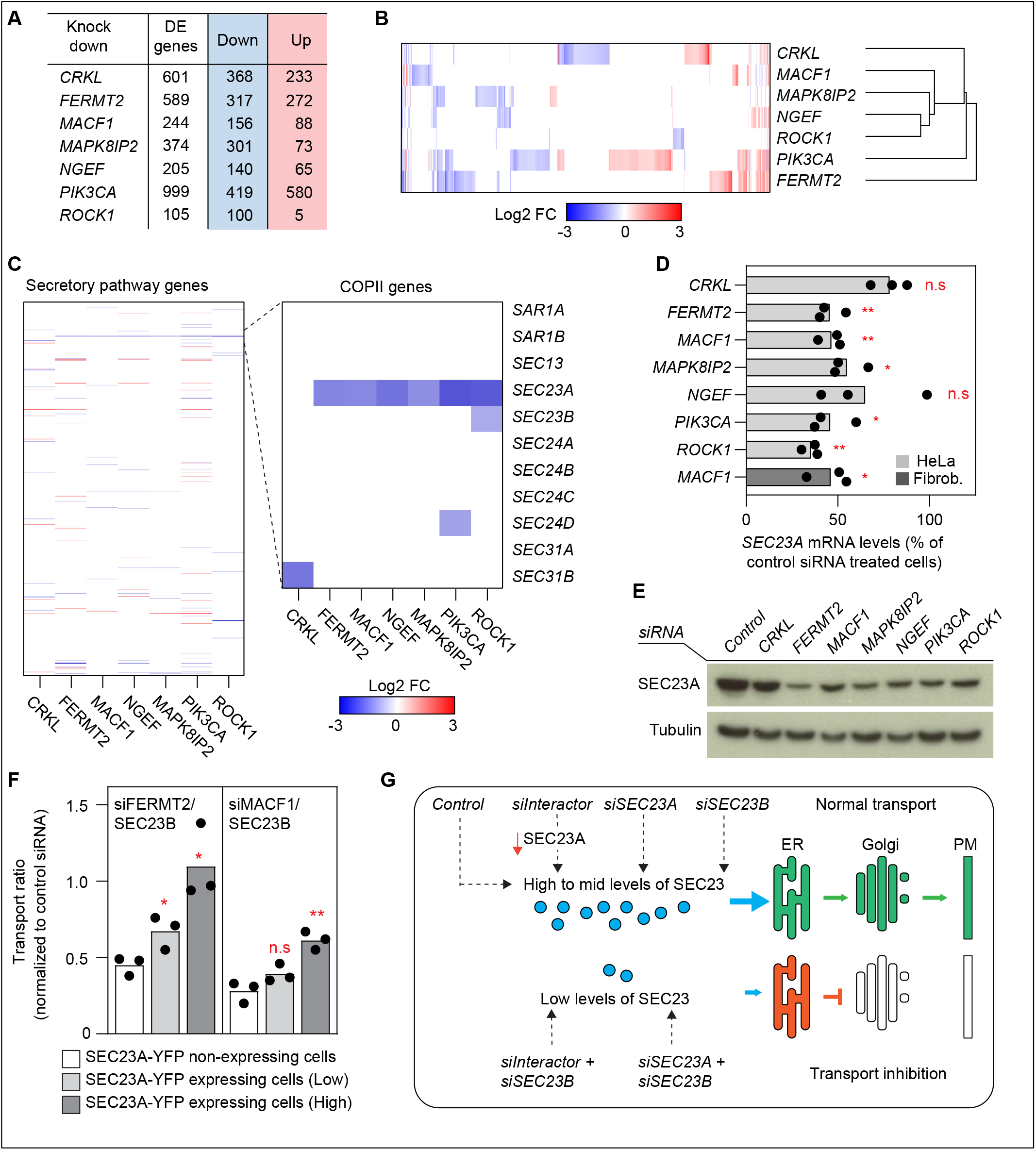
The transport defect phenotype is caused by downregulation of SEC23A. **(A)** Differential expression analysis. Cells were treated with the indicated siRNAs and after 48 h total RNA was extracted and mRNA was sequenced using an Illumina workflow. The differentially expressed genes were obtained comparing expression to control-treated cells with a minimum of 3 biological replicas per condition. The FDR was set at 0.1 and the Log2FC at 0.58. **(B)** The differential expression analysis was hierarchically clustered and represented as a heat map using Morpheus (https://software.broadinstitute.org/morpheus). **(C**) Analysis of a subset of 595 manually curated secretory pathway genes for the indicated knockdowns (48 h) compared to control siRNA. The zoomed-in region highlights the COPII genes. **(D)** RT-qPCR quantification of *SEC23A* levels after 48 h of knockdown with the indicated siRNAs. Bars represent averages of 3 independent experiments and the dots the individual values. The dark grey bar represents *SEC23A* levels after 48 h of MACF1 siRNA treatment in human lung fibroblasts. **(E)** Western-Blot analysis of SEC23A protein levels after 48 h of depletion with the indicated siRNAs. Alpha tubulin was used as a loading control. **(F)** SEC23A rescue experiment. Cells were co-transfected with the indicated siRNAs and a cDNA encoding SEC23A-YFP. After 60 h of transfection, the VSVG assay was performed. Based on the intensity of the YFP channel, cells were divided in *non-expressing*, *low-expressing*, and *high-expressing*. **(G)** Scheme representing the transport phenotypes obtained during the screen and their relationship to SEC23A and SEC23B levels. Statistical significance: * p<0.05 and ** p<0.01 compared to control, Student’s t-test.

To find a possible cause for the observed transport block when any of these interactors were doubled knocked-down with SEC23B, we started by focusing our analysis of the transcriptome data on a set of 595 secretory pathway-related genes based on a recently annotated gene list (Feizi et al. 2017). Overall, the transcriptional changes observed for those genes were relatively minor with no particular enrichment of these genes or parts of them in any condition (Fisher’s exact test). Surprisingly, we found that *SEC23A* was significantly downregulated in 6 of the 7 SEC23B functional interactors causing VSVG transport inhibition (**Fig. 3C** and **Table 5**). These 6 interactors are the same ones that showed the strong ER cargo retention in Fig 1D. Most remarkably, *SEC23A* was found to be the only common gene that was affected by knockdowns of all 6 interactors. The downregulation of *SEC23A* upon knockdown of the interactors could also be confirmed independently by RT-qPCR in HeLa cells and also in human lung fibroblasts for the case of *MACF1* (**Fig. 3D)**. The downregulation in *SEC23A* mRNA levels is also reflected at the protein level **(Fig. 3E)** and the transport phenotype can be rescued by restoring the levels of SEC23A with the overexpression of SEC23A-YFP in double knockdown cells (**Fig. 3F**). These results offer a straightforward explanation for the ER block in the double knockdown experiments described here, namely, reduction in the levels of total SEC23 by direct downregulation of *SEC23B* and an indirect downregulation of *SEC23A* (**Fig. 3G**). However, we cannot exclude the possibility that changes in other genes than *SEC23A* also contribute to the transport inhibition phenotype observed here.

### ECM related cues are connected to SEC23A expression

The decrease in SEC23A induced by the depletion of particular cytoskeleton regulatory and associated proteins, suggests that cytoskeleton-related cues are connected to SEC23A gene expression. But which are these cues?

Using Metascape (Zhou et al. 2019) we performed enrichment analysis on the mRNA-Seq data to look for the most relevant processes affected as a consequence of the depletion of these cytoskeleton-associated proteins that are connected to the expression of SEC23A. We found that the most common processes enriched among the analyzed proteins were cell adhesion, negative regulation of cell proliferation, and blood vessel development (**Fig. 4A** and **Table 6)**. We decided to focus on cell adhesion as it is a process that is directly connected to the cytoskeleton and some of the functional interactors described here have a well-established link to it. For instance, MACF1 has been associated with focal adhesions dynamics (Wu, Kodama, and Fuchs 2008) and FERMT2 regulates integrin-dependent cell-matrix attachment (Tu et al. 2003; Montanez et al. 2008). In fact, 4 out of these 6 functional interactors are bona fide members of the adhesome (Zaidel-Bar et al. 2007), which currently includes 232 proteins known to be an integral part of focal adhesions (FAs) or associated with them (www.adhesome.org) (**Fig. 4B**). Consistent with the idea that knockdown of these proteins induces changes in FAs, we observed morphological alterations in FAs as assessed by vinculin staining. For instance, we observed enlarged focal adhesions in the MACF1 knockdown cells, in agreement with a knockout mouse model described earlier (Wu, Kodama, and Fuchs 2008) and similar alterations were observed for the other interactors (**Fig. 4C**). These data suggest that changes in FAs or FA signalling might be connected to the expression of SEC23A. To test this, we treated cells with the FA kinase (FAK) inhibitor PND-1186. As seen in **Fig. 5A**, FAK inhibitor treatment upregulated specifically SEC23A but no SEC23B. Interestingly, FAK inhibitor treatment could only partially counteract the decrease in SEC23A levels induce by depletion of the interactors (**Fig. 5B**)

**Figure 4.**
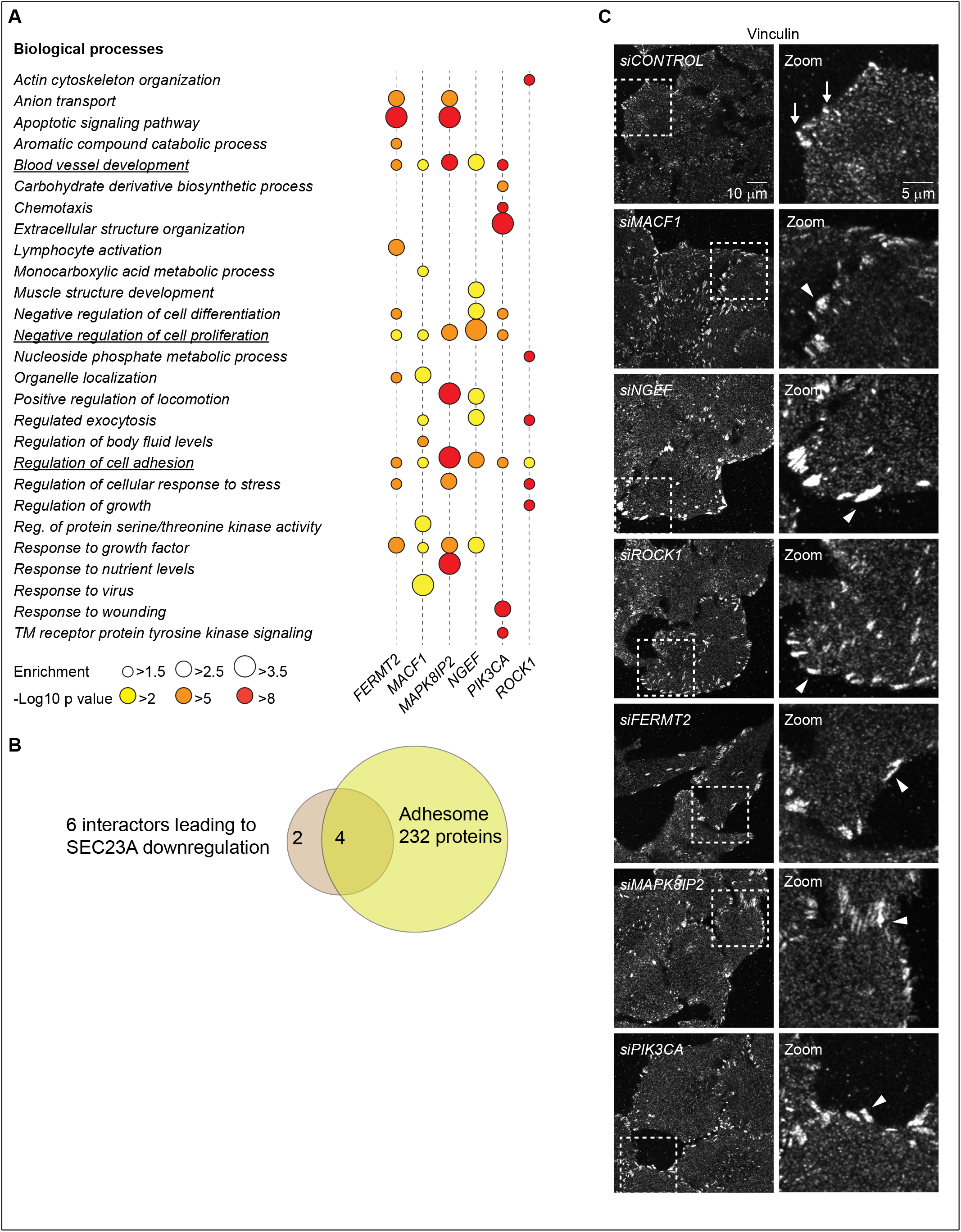
Knockdown of the SEC23 interactors causes changes in focal adhesions. (**A**) Gene enrichment analysis was done using Metascape. GO terms for biological processes with a minimum enrichment of 1.5 and a minimum p-value of 0.01 are depicted. When possible, child terms were used over parents and redundant pathways were omitted. The most common processes across all 6 conditions are underlined. **(B)** Venn diagram representing the intersection between the 6 interactors and the adhesome proteins. **(C)** Confocal images of focal adhesions assed by vinculin staining after 48 h of treatment with the indicated siRNAs. Arrows show the focal adhesions in control cells. Arrowheads show enlarged focal adhesions in knockdown cells.

**Figure 5.**
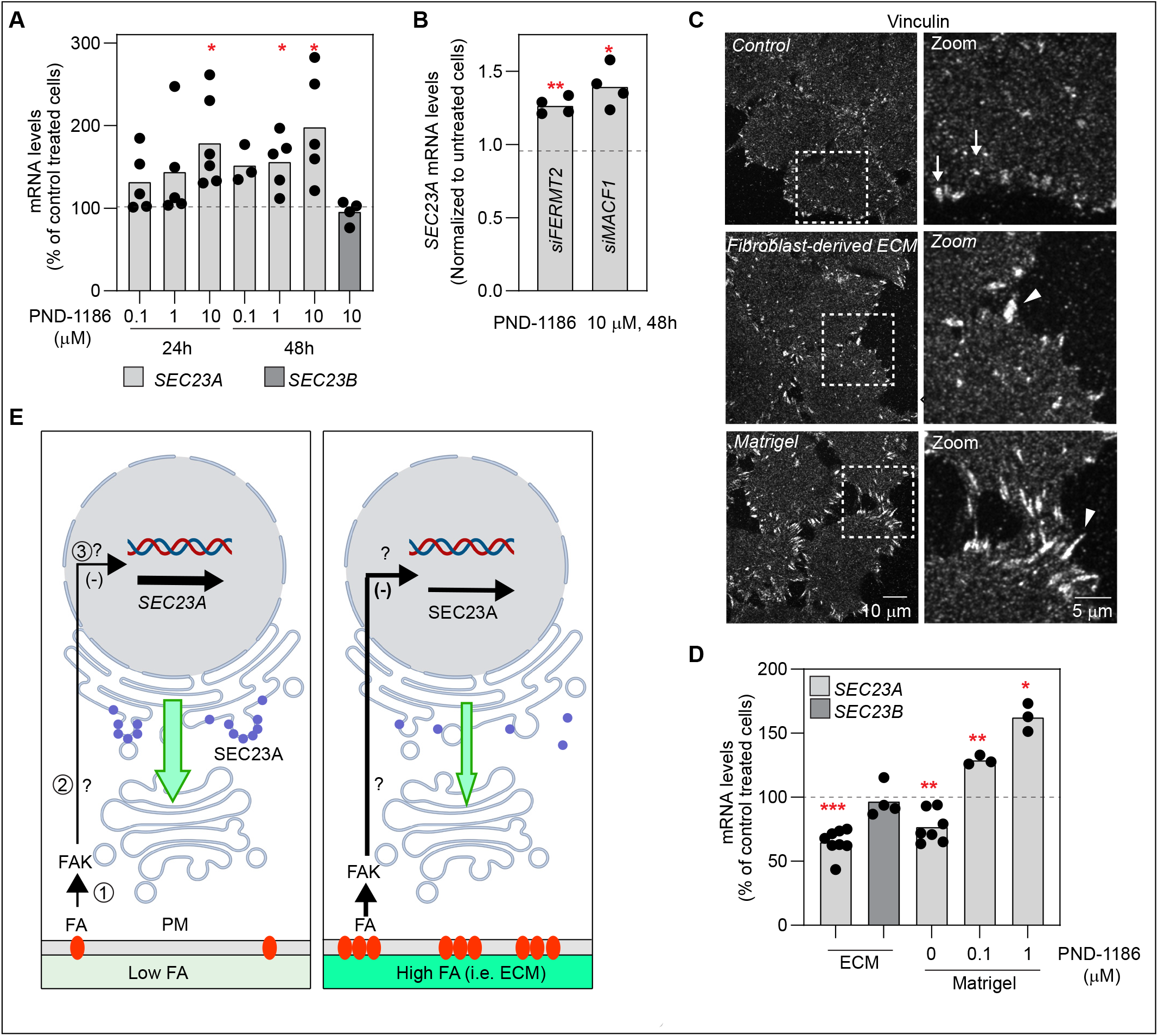
FAK signaling and cell attachment to ECM are connected to SEC23A expression. **(A)** *SEC23A* and *SEC23B* mRNA levels were assessed by RT-qPCR after treatment with the FAK inhibitor PND-1186 for the indicated times at the indicated concentrations. **(B)***SEC23A* mRNA levels were assessed by RT-qPCR in cells treated with the indicated siRNAs together with the FAK inhibitor at the indicated time and concentration. **(C)** Confocal images of FAs assessed by vinculin staining after 48 h of the indicated treatment. Arrows show the FAs in cells plated in control (plastic) plates. Arrowheads show the enlarged focal adhesions in cells plated in ECM or Matrigel treated plates. **(D)***SEC23A* and *SEC23B* mRNA levels were assessed by RT-qPCR after 48 h of seeding cells in dishes coated with Matrigel or fibroblast-derived ECM and treated with the FAK inhibitor PD-1186 for 24 h at the indicated concentrations. **(E)** Model representing the hypothetical relationship between FAs and SEC23A. (1) Interaction of cells with ECM or substrate through FA. (2) Activation of FAK and a downstream signaling cascade. (3) Transcriptional downregulation of SEC23A. (4) SEC23A-mediated ER to Golgi transport. In a condition of low FAs or low FAK signaling (left), *SEC23A* gene expression is not highly repressed. SEC23A levels are normal (blue circles) and SEC23A-dependent transport proceeds normally (green arrow). In a condition of high FA or high FAK signaling (right) the repression is more prominent and SEC23A levels decrease, possibly affecting SEC23A-dependent transport. The elements of the signaling cascade and the transcriptional regulation remain to be identified.

As changes in FAs are dependent on interactions with ECM (see review by (Geiger, Spatz, and Bershadsky 2009)) we hypothesized that SEC23A might be sensitive to this interaction. To test this more directly we plated cells specifically on different ECMs and measured the levels of *SEC23A* by RT-qPCR. Cells plated on Matrigel or in fibroblast derived-ECM showed reduced *SEC23A* mRNA levels compared to cells plated on plastic alone (**Fig. 5C**). Interestingly the Matrigel-mediated downregulation of *SEC23A* could be prevented by treatment with the focal adhesion kinase (FAK) inhibitor PND-1186 (**Fig. 5D**).

## DISCUSSION

In this work, we have studied the potential functional interaction of 378 cytoskeleton-related proteins with the COPII component SEC23 in the early secretory pathway. This uncovered, amongst others, six strong SEC23 functional interactors that have been proposed to be involved in cell attachment to ECM through FAs. Surprisingly, single knockdowns of each of these proteins did not only lead to morphological changes in FAs and changes in genes connected to adhesion but also to the transcriptional down-regulation of *SEC23A*. While these results provide a plausible explanation of our screening results showing that co-downregulation of any of these proteins together with SEC23B causes a strong transport inhibition at the ER level, most likely due to a strong depletion of both SEC23 paralogues, they also connect for the first time FAs and ECM-related cues to the COPII machinery. Independent experiments here, in support of this hypothesis, showed, for instance, that plating cells on fibroblast-derived ECM or Matrigel led to the downregulation of *SEC23A*, an effect that is dependent on FAK activity. Collectively, these results suggest that cell attachment to ECM can regulate the levels of SEC23A in a process that involves FA signalling. These ideas are summarized in the model shown in Fig. 5E. We propose that FA signaling exerts a negative regulation of SEC23A levels. In the context of low FA or low FAK signaling (i.e cells plated on plastic), the levels of SEC23A remain relatively constant. In the context of high FA or FAK signaling (i.e cells plated on ECM), SEC23A is downregulated. These results contribute to understanding how cells can integrate extracellular information such as ECM signaling into the secretory pathway and expands the increasing set of cues that are proposed to regulate the early secretory pathway including nutrient sensing (Zacharogianni et al. 2011; Liu et al. 2019) and growth factor signaling (Farhan et al. 2010; Tillmann et al. 2015; Scharaw et al. 2016).

Our experimental approach consisted of an unbiased siRNA-based double knockdown screen. This kind of experimental strategy has been used by others to identify complex networks underlying, for instance, cholesterol metabolism (Zimon et al., 2020 https://doi.org/10.1101/2020.10.29.360818), cancer (Laufer et al. 2013), or chromatin remodelling (Roguev et al. 2013). Here, this strategy led us to the identification of 20 cytoskeleton-related proteins that functionally interact strongly with either SEC23A or SEC23B in the transport of the well-characterized cargo model VSVG. The majority of these 20 proteins have not been previously implicated in the secretory pathway. Furthermore, none of them were identified as hits in an earlier genome-wide siRNA screen which identified several hundred human genes directly or indirectly involved in biosynthetic transport from the ER to the plasma membrane (Simpson et al. 2012) or in another genome wide screen in Drosophila (Bard et al. 2006). Altogether, this demonstrates the potential of such double knockdown functional interaction screens as a tool to identify new regulators of the processes under view.

Previous works have demonstrated a significant role of the cytoskeleton as a facilitator of biosynthetic membrane traffic in the early secretory pathway including microtubules and associated microtubule motor proteins (Scales, Pepperkok, and Kreis 1997; Presley et al. 1997; Watson et al. 2005; Brown, Hunt, and Stephens 2014; Verissimo et al. 2015). Therefore, we expected to find in our screen here many proteins with a more canonical role in the cytoskeleton, such as motor or alike proteins. While a few of such proteins were indeed identified in this screen, the majority of the proteins that we ranked as the 20 strongest functional interactors of SEC23 in our VSVG transport assay are described as regulators of signalling pathways at the plasma membrane/ECM interface.

Interestingly, our results here show an apparent lack of overlap between SEC23A and SEC23B functional interactors. Although these two SEC23 paralogues are 85% identical at the amino acid sequence level, they have been proposed to have different functions in terms of cargo specificity. SEC23A has been proposed to mediate transport of extracellular matrix components, specifically collagens (Boyadjiev et al. 2006; Lang et al. 2006), and SEC23B has been involved in the maintenance of professional secretory tissues like the pancreas (Khoriaty et al. 2016, 2017) or to the transport of EGFR (Scharaw et al. 2016). However, this view has been recently challenged by demonstrating an indistinguishable protein interactome of the two paralogues and the possibility to exchange the protein domains amongst them without apparent functional losses (Khoriaty et al. 2018). It was proposed instead that the SEC23 paralogues may underlie different transcriptional regulations which could in part explain the distinct phenotypes of SEC23A/B deficiency within and across different species (Khoriaty et al. 2018). Our results here support these findings as, first, the knockdown of SEC23A or SEC23B alone did not result in a significant transport inhibition of VSVG, suggesting that the remaining paralogue can replace at least partially the function of its counterpart under the conditions and transport marker used here. And second, while the transcriptional regulation of SEC23B is proposed to be connected to growth factor signaling (Scharaw et al. 2016), here we have shown that transcriptional regulation of SEC23A appears to be linked to ECM signaling, a scenario which would explain the lack of common functional interactors in our screen.

Within the 20 interactors found in the screen, we focused on a subgroup that led to a strong ER block phenotype (FERMT2, MACF1, MAPK8IP2, NGEF, PIK3CA, ROCK1). Most of these proteins are involved in cell attachment to ECM through FAs. For instance, FERMT2 is located at FAs and participates in the connection of ECM to the actin cytoskeleton through integrins (Tu et al. 2003), MACF1 crosslinks actin and microtubules at FAs (Wu, Kodama, and Fuchs 2008), ROCK1 is involved in stress fibre and FAs formation (Amano et al. 1997) and PIK3CA is a bona fide member of the FA interactome (www.adhesome.org). The role of FAs in the interpretation of ECM-related cues is well established (Geiger, Spatz, and Bershadsky 2009). FAs have also been involved in membrane trafficking including endocytosis and exocytosis (Wickström and Fässler 2011; Nolte, Hoen, and Margadant 2020). In exocytosis, FAs appear to serve as hot spots for secretion, a function that might be linked to the role of FAs in the organization of microtubules from TGN to the PM (Stehbens et al. 2014; Fourriere et al. 2020; Eisler et al. 2018). Based on our results here, we propose that ECM and FAs are also connected to membrane trafficking through the regulation of SEC23A expression. These views are complementary and contribute to understanding how cells can connect trafficking and extracellular cues in a robust manner. Since SEC23A has been proposed to mediate transport of ECM components, specifically collagens (Boyadjiev et al. 2006; Lang et al. 2006) linking the ECM to the expression of SEC23A at the transcriptional level, as suggested by our results here, offers a potentially interesting feedback mechanism in which signals originating at the ECM could, in turn, regulate the deposition of ECM components by controlling the levels of SEC23A. Finally, the ECM-mediated regulation of SEC23A might also be relevant for cancer. It has been shown that downregulation of SEC23A leads to changes in the secretome of cancer cells that promote metastatic capabilities (Korpal et al. 2011; Szczyrba et al. 2011). As metastatic cancer cells migrate and interact dynamically with the ECM it is plausible that signals arising from the ECM contribute to SEC23A regulation in cancer cells, although this needs further investigation. The precise molecular cascade that connects ECM to the expression of SEC23A and its physiological implications remain to be investigated.

## MATERIALS AND METHODS

### Cell lines and reagents

HeLa Kyoto cells were a gift from Shuh Narumiya (Kyoto University) and were cultured in DMEM supplemented with 10% FCS and 2 mM glutamine. Normal human primary lung fibroblasts were obtained from Lonza (Basel, Switzerland) and were cultured in fibroblast growth basal medium. HeLa cells stably expressing P24-YFP (TMED2) were generated in-house and describe before (Simpson, Nilsson, and Pepperkok 2006). All cells were kept in a humidified incubator at 37°C and 5% CO_2_.

All siRNAs used were from Ambion and are described in **Table 1**. Anti VSVG antibody and recombinant adenovirus expressing YFP-tagged and CFP-tagged tsO45G thermosensitive versions of VSVG were a gift from Kai Simmons (MPI-CBG, Dresden, Germany). Large-scale adenovirus preparation was done by Vector Biolabs. RUSH construct encoding E-Cadherin-GFP was a gift from Frank Perez (Institute Curie, Paris, France). YFP–SEC23A was described elsewhere (Stephens et al. 2000). Anti MACF1 antibody was a gift from Ronald Liem (Columbia University, NY, USA). The following primary antibodies were used: SEC23A (ab179811) from Abcam, Vinculin (MAB3574) from Merck, EEA1 (610457) and GM130 (610822) from BD biosciences, Alpha-Tubulin (MS581P1) from Thermo Fisher, SEC23B (GTX82432) from GeneTex, GFP (11814460001) from Roche. Anti-beta COPI rabbit serum was produced in-house. The following secondary antibodies were used: rabbit anti-mouse-HRP (A9044) from Merck, goat anti-rabbit-HRP (ab6721) from Abcam and Alexa-conjugated antibodies from Molecular probes. Hoechst dye 33258 was from Invitrogen. SiR-DNA was from Spirochrome. FAK inhibitor PND-1186 was from Tocris Bioscience. Matrigel was from Corning. Cycloheximide was from Calbiochem, Biotin was from Sigma.

### siRNA-based functional interaction screen

The cytoskeleton siRNA library was designed in-house and contains 848 siRNAs targeting 378 cytoskeleton, associated, and regulatory proteins (**Table 1**). The cytoskeleton siRNAs were spotted on Nunc Lab-Tek chambers at a density of 384 spots per chamber as previously described (Simpson et al. 2012). The whole cytoskeleton library was distributed on 4 Lab-Tek chambers which included also positive controls (beta and gamma *COPI* targeting siRNAs), 2 different non-targeting siRNAs (Scramble and Neg9 siRNAs), and an *INCENP* targeting siRNA to evaluate the transfection efficiency of the plate based on the number of multinucleated cells (Neumann et al. 2006). Spotted Lab-Teks were dried, stored in humidity-free containers, and used within a month after coating. At the moment of cell plating, cells were transfected either with *SEC23A*, *SEC23B*, or control siRNAs in suspension at a final concentration of 15 nM using Lipofectamine 2000 and Opti-MEM (Thermo Fisher) according to the manufacturer’s instructions. 1.5 mL of transfected cell suspension containing 100,000 cells were plated onto each Lab-Tek. At 48 hrs after plating, cells were infected with adenovirus coding for the VSVG-tsO45-YFP protein (henceforth called VSVG-YFP) for 1 hr at 37°C, washed twice and transferred to 40°C for 12 hrs to accumulate VSVG-YFP in the ER as described (see Simpson et al., 2012). Release of the transport marker from the ER was induced by shifting temperature to 32°C in fresh DMEM containing 25 mM HEPES and 50 μg/mL of cycloheximide. After 1 hr at 32°C cells were washed twice with PBS and fixed with PFA 3% for 20 min. VSVG at the plasma membrane was detected by incubating cells for 1h with an anti-VSVG antibody (1:100) recognizing an extracellular epitope of VSVG, followed by 30 min incubation of anti-mouse Alexa-647 secondary antibody (1:500). Cell nuclei were stained with Hoechst dye 33258. Finally, cells were washed twice with PBS and left with PBS for imaging. Images were acquired on an Olympus ScanR microscope using a 20X/0.7 NA objective. The whole screen was carried out in 5 independent biological replicas.

### Screen data analysis

Images were analyzed using CellProfiler (Carpenter et al. 2006) as previously described (Scharaw et al. 2016). For the analysis, only images that contained between 5-200 cells were considered. The transport ratio per cell was defined as the VSVG specific intensity at the plasma membrane (A647 signal) divided by the VSVG-YFP intensity (representing the total cellular amount of VSVG expressed). Transport ratios for each siRNA were converted to transport scores according to the following formula: Transport score = *xi−X/MAD*, where xi is the average transport ratio per cell for the siRNA of interest, X is determined as the median of 25 (5×5 matrix) transport ratios from neighbouring siRNA sports surrounding the siRNA spot of interest. MAD is the median absolute deviation in the 5×5 matrix. (**Table 2**).

Based on our positive controls (*COPB1, COPG1, SEC23A/B*) we set our threshold on a transport score of −1.5 (strong inhibition). By symmetry, only combinations showing scores above 1.5 were considered as transport accelerators. Both non-targeting siRNAs (Scramble and Neg9) had transport scores close to 0 (0.08 and −0.16 respectively) so we selected Neg9 as a control siRNA for our following analysis. As a quality control, we only considered the plates in which our positive controls showed strong transport inhibition and the negatives controls had a score close very close to 0. Finally, we considered only the siRNAs for which we had at least 3 biological replicas and we computed the median transport score of these replicas for each siRNA. To obtain the hits, we compared the added median transport scores of the single knockdowns vs the double knockdowns and selected the ones in which the difference was greater than 1 transport score unit (**Table 2** and Table legends).

### Confocal microscopy and image segmentation

Confocal stacks were acquired using a Leica SP8 system with a 63X/1.4 NA objective and Zeiss 780 with a 40X/1.1 NA objective. Unless otherwise stated, the images shown represent maximum intensity Z-projections. Pipelines for quantification of ERES and COPI transport intermediates were implemented as custom Fiji plugins. ERES and COPI structures were segmented and quantified in 3D in the individual cells using functions of the 3D image suite library (Ollion et al. 2013). First nuclei were segmented in the Hoechst channel. Cell masks were identified by applying a watershed algorithm in the COPI channel using previously segmented nuclei as seeds. For robust segmentation of ERES and COPI positive structures, 3D median filters were applied for noise suppression and for subtracting local backgrounds in corresponding channels. Seeds for the potential structures were identified by applying a local maximum filter. Individual structures were segmented using spot segmentation plugin of 3D image suite including watershed separation of touching structures. When processing the COPI channel, big positive clusters, which presumably correspond to the Golgi complex, were segmented in 3D by thresholding and excluded from spot analysis. The pipeline documented the number of identified structures in each cell, their integrated intensity, and the integrated intensity of the entire cell.

### RUSH assay

For the RUSH assay the cells were plated in coverslips in 24-well plates at a density of 15,000 cells per well and 24 hrs later transfected with siRNAs using Lipofectamine 2000 and serum-free DMEM. 24 hrs later the cells were transfected with the RUSH construct encoding E-cadherin using Lipofectamine LTX and Enhancer in serum-free DMEM (as per manufacturer’s instructions). The next day the RUSH assay was performed as described previously (Boncompain et al. 2012). In brief, the cell medium was changed to fresh DMEM containing 40 μM biotin and 50 μg/mL of cycloheximide. After 45 min cells were washed once with ice-cold PBS and incubated with anti-GFP (1:100) in PBS on ice for 45 min. After incubation, cells were washed twice with PBS and fixed with PFA 3% at room temperature for 20 min. After washing two times with PBS cells were stained for 30 min with an anti-mouse Alexa-647 antibody (1:500). Cell nuclei were stained with Hoechst dye 33258. Finally, coverslips were mounted on glass slides using Mowiol mounting media, and images were acquired and analyzed with CellProfiler as above. For every condition, a non-release control was also included to account for cargo leakiness.

### Electron microscopy

Cell monolayers were fixed in 2.5% glutaraldehyde in cacodylate buffer, washed with cacodylate, and post-fixed in 1% osmium. En-bloc staining was done in 1% uranyl acetate (UA). The cells were then dehydrated in ethanol, and embedded in Epon resin. All specimen preparation steps up to embedding were done using a PELCO Biowave Pro microwave (Lorentzen et al. 2018). Thin sections of 70 nm were collected in 2×1 slot grids and post-stained with lead citrate. The imaging was done using LLP viewer in a JEOL 2200 Plus electron microscope equipped with a Matataki camera.

### Co-immunoprecipitation and Western Blot analysis

For Co-IP experiments, Hela cells were plated on a 10 cm Petri dish and lysed at approximately 90% of confluency using NP-40 lysis buffer (1% NP-40, 150 mM NaCl, 50 mM Tris/HCl pH 7.4) supplemented with protease inhibitors. All subsequent steps were carried out at 4°C. Cells were lysed for 30 min and centrifuged at 20,000 g for 15 min. 60 μL of protein G-agarose were added to 400 μL of lysate in the presence or absence of 2 μg of anti-MACF1 antibody and incubated with rotation overnight. Beads were pelleted and washed 3 times with lysis buffer and then heated at 60°C for 5 min with sample buffer 2x (10% glycerol, 4.5% SDS, 130 mM DDT, 0.005% bromophenol blue, 80 mM Tris/HCl pH 6.8). Proteins were separated by SDS-PAGE using a 3-8% Tris-Acetate gel and then transferred to PVDF membranes for Western blotting.

For SEC23A experiments, cells were plated in 12 well plates, treated with the respective siRNAs for 48 h, and then lysed directly with sample buffer 2x. Lysates were treated with Benzonase for 10 min at room temperature and then heated at 90°C for 5 min. Proteins were separated by SDS-PAGE using a 4-12% Bis-Tris gel and transferred to PVDF membranes as above.

### mRNA-Seq

Individually barcoded stranded mRNA-Seq libraries were prepared from high-quality total RNA samples (~600 ng/sample) using the Illumina TruSeq RNA Sample Preparation v2 Kit (Illumina) implemented on the liquid handling robot Beckman FXP2. Obtained libraries that passed the QC step were pooled in equimolar amounts; 1.8 pM solution of this pool was loaded on the Illumina sequencer NextSeq 500 and sequenced uni-directionally, generating ~500 million reads, each 85 bases long.

Single-end sequencing reads were aligned to the human genome (version GRCh38) and the reference gene annotation (release 84, Ensembl) using STAR v2.6.0a (Dobin et al. 2013) with default parameters. Read counts per gene matrices were generated during the alignment step (--quantMode GeneCounts). Gene counts were then compared between 2 groups of replicated samples using the DESeq R package (Love, Huber, and Anders 2014). Differentially expressed genes were selected based on their p-adjusted value lower than 10% and their Log2 fold change above 0.58 or below −0.58

### RT-qPCR

Total RNA was extracted using RNAeasy kit (Qiagen). 500 ng of total RNA was subjected to reverse transcription using SuperScript III First-Strand Synthesis Supermix (Invitrogen) according to the manufacturer’s instructions. cDNAs obtained this way were diluted 1/10 and used for PCR. The RT-qPCR reaction was performed using the SYBR green detection reagent (Applied Biosystems) in StepOne Real-Time PCR System machines using the StepOne software v2.3 (Applied Biosystems). Primer sequences are described in the supplementary information. Changes in gene expression were calculated with the 2^−ΔΔCT^ method (Livak and Schmittgen 2001) using *GAPDH* as the housekeeping gene.

### SEC23A rescue experiment

Cells plated in 24 glass-bottom well plates were co-transfected with siRNAs and SEC23A-YFP using Lipofectamine 2000. After 48 h cells were infected with VSVG-CFP-coding adenovirus and the VSVG assay was carried out as previously indicated with the following modifications to minimize fluorescence bleed-through: the external VSVG was detected using a secondary Alexa-568-conjugated antibody and the nuclei were stained in the far red using SiR-DNA. The transport ratios were obtained as previously indicated and cells were separated into 3 categories based on the intensity in the YFP channel: non expressing cells, low expressing cells, and high expressing cells.

### ECM experiments

For preparing ECM derived from lung fibroblasts, cells were seeded at a density of 20,000-25,000 cells/well in 24 glass-bottom well plates. After 18 hrs cells were switched to DMEM medium with macromolecular crowding (a mixture of Ficol 70 and Ficol 400) (Chen et al. 2009). Cells were cultured for 5 days with stimulation (TGFß1 5 ng/ml) and media change on day 1 and day 4. On day 5, cells were washed with PBS and the plates were subjected to decellularization using 1% NP-40 (Harris, Raitman, and Schwarzbauer 2018). For Matrigel experiments, Matrigel was diluted in DMEM at a final concentration of 0.2 mg/ml and used to cover plates according to the manufacturer’s instructions.

## SUPPLEMENTARY FIGURE LEGEND

**Figure S1.**
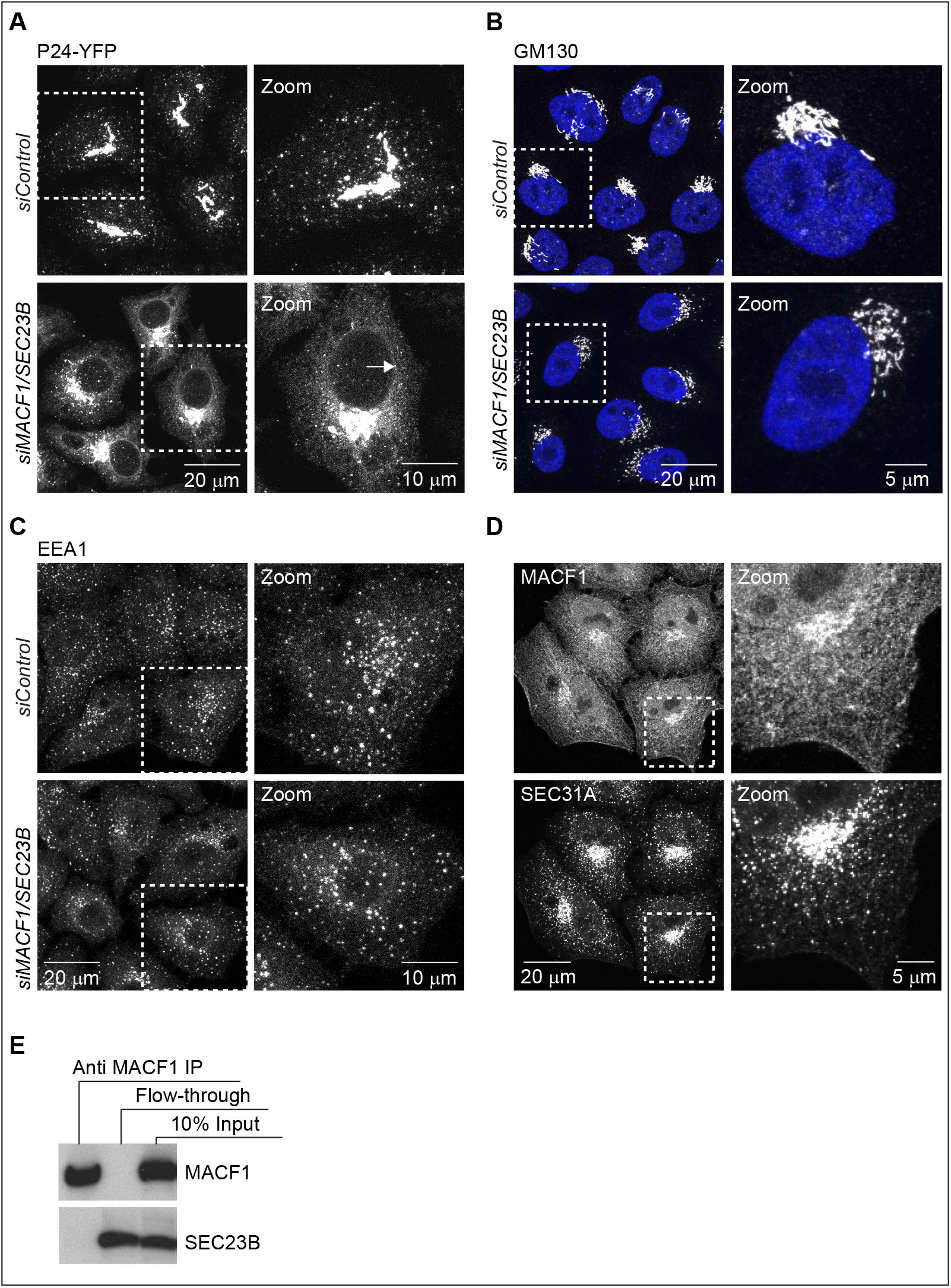
MACF1/SEC23 double knockdown affects ER exit. **(A)** Confocal images of HeLa cells stably expressing P24-YFP and transfected with the indicated siRNAs for 48 h. Arrows indicate ER membranes. Confocal images of cells transfected with the indicated siRNAs for 48 h, fixed and stained with **(B)** anti GM130 antibody or **(C)** anti EEA1 antibody. **(D)** Confocal images of untreated HeLa cells fixed and co-stained for MACF1 and SEC31A as a marker for ERES. **(E)** Co-IP experiment. HeLa cells were lysed and incubated with anti MACF1 antibody bound to agarose-protein G beads at 4°C overnight. The beads were washed and loaded into gels. Western-Blot was performed using anti MACF1 and anti SEC23B antibodies.

## TABLE INFORMATION

**Table 1. Cytoskeleton siRNA library and other siRNAs**

All siRNA used are predesigned Silencer Select from Ambion except otherwise stated. They were selected based on the score and number of transcripts targeted.

**Table 2. Screen results**

The first column, “spotted siRNA”, refers to the cytoskeleton library. The second column refers to the siRNAs transfected in liquid suspension. The third column shows the median value of all the biological replicas for a given combination. The results shown here include only the plates that passed quality control.

**Table 3. SEC23 interactors**

The top 25 double knockdown combinations are listed according to transport scores. The expected additive effects refer to the expected transport score if the siRNA perturbations would have an additive effect. (Observed – expected) refers to the transport score difference between the observed score in the double knockdown minus the expected score assuming additive effect.

The flagged combinations were not considered. The flags are the following: 1) Value in single knockdown is too high (>0.75 or <-0.75), 2) Difference between observed and expected transport scores is less than 1 transport score unit, 3) Gene not expressed in HeLa cells (according to our transcriptomics data), 4) Less than 3 biological replicas, 5) Transport score in double knockdown below the 1.5 threshold, 6) siRNA does not target the specified gene in current Ensembl version GRCh38.

**Table 4. Differential expression analysis**

The differential expression analysis was set at −0.58>Log2FC >0.58 and the p-adjusted at 0.1. The values listed correspond to individual knockdowns compared to the control non-targeting siRNA

**Table 5. Secretory pathway related genes**

Differential expression analysis for a set of 595 annotated and manually curated secretory pathway genes based on a list previously published (Feizi et al. 2017)

**Table 6. Gene enrichment**

Top 50 GO: biological processes ranked by the number of genes for each knockdown. The enrichment value was set at a minimum of 1.5 and the p-value at 0.01.

## ACKNOWLEDGMENTS

We would like to thank the EMBL Advanced Light Microscopy Facility for help with the preparation and analysis of the screen, the EMBL Genomics Core Facility for help with mRNA-Seq sample preparation and analysis, Seetharaman Parashuraman (Institute of Biochemistry and Cell Biology, Naples, Italy) for useful discussions and critical reading of the manuscript, Charlotte Carr for help with the RUSH assay, and members of the Pepperkok Team for fruitful discussions during the development of this project. JJ was funded by a Fellowship from National Commission for Scientific and Technological Research, Chile. MMK was funded by a BMBF grant (German Centre for Lung Research, DZL).

## AUTHOR CONTRIBUTION

Conceptualization: Juan Jung, Rainer Pepperkok, Muzamil M. Khan

Investigation: Juan Jung, Muzamil M. Khan

Formal Analysis: Juan Jung, Muzamil M. Khan, Jonathan Landry, Aliaksandr Halavatyi

Resources: Pedro Machado, Miriam Reiss

Supervision: Rainer Pepperkok

Writing original draft: Juan Jung, Rainer Pepperkok, Muzamil M. Khan

Writing review and editing: all the authors

Funding acquisition Rainer Pepperkok, Juan Jung

